# The Mismatch Negativity compared: EEG, SQUID-MEG and novel ^4^Helium-OPMs

**DOI:** 10.1101/2025.04.04.647154

**Authors:** Tjerk P. Gutteling, Jérémie Mattout, Sébastien Daligault, Julien Jung, Etienne Labyt, Denis Schwartz, Françoise Lecaignard

## Abstract

Magneto-encephalography (MEG) provides a higher spatial resolution than electro-encephalography (EEG) to measure human auditory responses. However, conventional cryogenic MEG systems (SQUID-MEG) suffer from severe technological restrictions limiting, for instance, routine clinical use. Fortunately, a new generation of MEG sensors, optically pumped magnetometers (OPMs), have been developed to bridge the gap, combining the wearability of EEG with the benefits of MEG signal acquisition. We aim to assess their potential for studying auditory mismatch processing. The auditory Mismatch Negativity (MMN) is a well-characterized evoked component observable using a passive oddball paradigm with two-tone sound sequences. It has been extensively described using both EEG and MEG and is part of many EEG-based clinical applications, such as the assessment of patients with disorders of consciousness. MMN is therefore a relevant candidate to evaluate OPM performance. We use recently developed Helium-OPMs, which are high dynamic range MEG sensors that operate at room temperature. We compare their performance with cryogenic SQUID-MEG and EEG in a passive frequency oddball paradigm. Results show a significant MMN across subjects in all modalities as well as a high temporal similarity between modalities. Signal-to-noise ratios were also similar, and detection of significant individual MMN (within-subjects) using the OPM system was equal or better than EEG. Given that the OPM system tested here is a prototype comprised of only five sensors, these results are a promising step towards wearable MEG that combines the advantages of MEG and EEG.

## Introduction

The Mismatch Negativity (MMN) is simple to evoke, yet complex to unravel. Auditory MMN is one of the most extensively studied neural biomarkers of sensory processing, and led to in-depth knowledge of perception as an inference process (Fitzgerald and Todd, 2020; Näätänen et al., 2007). This electrophysiological component, first described in Näätänen et al. (1978), is typically observed using auditory oddball paradigms, where an ordered sequence of stimuli (typically repeated tones, called standards) is interrupted by infrequent different stimuli (called deviants). MMN is observed by subtracting the standard evoked response from the deviant one, and generally peaks between 100ms and 250ms after sound onset. Key aspects of the MMN include its automaticity (it can be measured without paying attention to the sounds) and its persistence in sleep and states of unconsciousness (Morlet and Fischer, 2014). Not only has MMN been invaluable to study sensory processing with high temporal resolution but it has become an essential tool in cognitive neurophysiology research (for review, see Csépe and Honbolygó, 2024; Näätänen et al., 2011). Its relevance endures today, as its widely accepted interpretation as a prediction error motivates its use to study the mechanistic foundations of predictive coding (Heilbron and Chait, 2018).

It is therefore not surprising that MMN has also proven to be highly informative in characterizing a variety of clinical disorders (Näätänen, 2003). The simplicity of the experimental set-up, which can be achieved without specific instructions, makes it an ideal tool for highly impaired clinical populations, as well as for infants. MMN could reveal a better understanding of the pathophysiology of several disorders, such as schizophrenia (Light and Swerdlow, 2015; Michie et al., 2016; Näätänen et al., 2016; Shelley et al., 1991), vegetative state outlook (Kane, 1996; Morlet et al., 2023), autism (Kujala et al., 2010), ADHD (Gomes et al., 2013), the assessment of preserved cognitive functions in comatose patients (Fischer et al., 2010; Morlet and Fischer, 2014) and more (for review see Schall, 2016). Furthermore, MMN can be used as a tool in brain computer interface (BCI) applications (Jin et al., 2015; Séguin et al., 2024).

Currently, electro-encephalography (EEG) is the standard for recording the MMN, especially in clinical populations, as it is portable, fairly cheap, robust to noisy environments and can be adapted to suit a wide range of recording conditions. While previous research has mostly investigated different parameters affecting the MMN (e.g. tone frequency, attentional modulation, statistical structure of the oddball stream), the focus is shifting towards more mechanistic research to unravel the still debated role of the MMN (Dürschmid et al., 2016; Lecaignard et al., 2022; Maheu et al., 2019) as well as its underlying physiology with finer spatial scale (Garrido et al., 2009; Lecaignard et al., 2021; López-Caballero et al., 2024; Valt et al., 2024). Here, magneto-encephalography (MEG) has its advantages due to the higher spatial resolution and better signal-to-noise ratio (Lecaignard et al., 2021). Where the volume currents that are picked up by the EEG are distorted by conductive head tissue, the magnetic signal picked up with MEG can pass relatively unhindered, giving it a spatial precision advantage as well as reduced interference from muscle artefacts (Claus et al., 2012; Gross, 2019). This, however, comes at the cost of severe restrictions, as the current MEG systems rely on superconducting quantum interference devices (SQUIDs, Cohen 1972) which require liquid helium cooling to operate and are therefore housed in a large rigid dewar. This implies that the classic MEG system is non-portable, and requires participants (patients) to come to the MEG site, which may be difficult for a clinical population. The sensor array is fixed and at a distance from the scalp due to an insulation layer, and participants are required to minimize any movement to prevent muscle artifacts and brain-to-sensor variability (Lopes da Silva, 2013; Puce and Hämäläinen, 2017). Furthermore, due to the necessity of a shielded room and helium cooling, cryogenic MEG costs vastly exceed those of EEG.

Fortunately, optically pumped magnetometers (OPMs) have recently been developed that provide the benefits of MEG with the increased mobility and flexibility of EEG (Brookes et al., 2022), although OPMs still require some form of shielding to operate correctly. OPMs are wearable MEG sensors that use density fluctuations of a gas cell probed by a laser to record variations in the magnetic field. As with EEG, the sensor can be mounted on a headcap or flexible helmet which can conform to any head shape and size. This is a benefit over cryogenic MEG, especially for younger populations with smaller head sizes, as it provides more consistent signal strength across the scalp. Due to the rigid helmets used with SQUID-MEG there is often an uneven signal quality, with a reduction of signal in the frontal and lateral areas. This is an important benefit given that the main cortical generators of MMN have been found in frontal and temporal areas (Deouell, 2008; Giard et al., 1995; Lecaignard et al., 2021; Recasens et al., 2015; Schönwiesner et al., 2007).

The majority of OPMs currently in use employ an alkali gas (usually Rubidium). These sensors usually have excellent sensitivity (7-15 ft/√Hz, compared to 3-4 ft/√Hz for SQUID-MEG), but suffer from a limited dynamic range (∼± 5 nT) (Allred et al., 2002; Tierney et al., 2019), which means that sensors can easily go outside their useable range due to head movement or fluctuations in the (external) magnetic field, causing signal loss. Contrary to SQUID-MEG, these sensors need heating to ∼150°C to operate, necessitating some insulation between the sensor and scalp. Here we use recently developed Helium-4 gas OPMs (^4^He-OPMs, MAG4Health, Grenoble, France). These sensors are natively triaxial, measuring signals in both the radial and tangential directions, operate at room temperature and have a much larger dynamic range (> 200 nT) (Beato et al., 2018; Labyt et al., 2019). The latter enables data acquisition with only modest magnetic shielding, such as a minimally shielded room in a hospital. This comes at the cost of some sensitivity, which, at time of data acquisition, is around 45 ft/√Hz on two of the three axes (Fourcault et al., 2021). Since they operate at room temperature and thus need no insulation, these sensors can be placed a little closer to the head, somewhat mitigating the loss of sensitivity (Bonnet et al., 2025; Iivanainen et al., 2017). In a previous study using visual and somatosensory stimulation, we showed that ^4^He-OPMs can provide very similar event-related fields (ERFs) and signal-to-noise ratios compared to SQUID-MEG (Gutteling et al., 2023).

Here we evaluate the performance of a prototype, 5-sensor ^4^He-OPM system in comparison with SQUID-MEG in the context of auditory mismatch processing. Given the importance of MMN in clinics using EEG, we also aim at comparing ^4^He-OPMs with EEG signals. Based on our previous evaluation of these sensors, we expect the resulting MMN as captured by the OPM system to be comparable to those obtained using EEG or SQUID-MEG. We will also explore the differences in ERFs between the radial and tangential axis of the OPM sensors. MEG is less sensitive to (quasi-)radially oriented sources than EEG due to the lack of scalp magnetic field produced by these sources. The additional tangential measurement axis of the OPM sensors may provide a more complete description of the magnetic field, potentially enabling a better characterization of radial sources unseen by cryogenic radial-only MEG (Haueisen et al., 2012).

## Materials and methods

### Participants

A total of 17 healthy participants (age range 19-51, mean age 31.6 ±8.5 years) took part in the current experiment as part of a larger study with multiple tasks (Gutteling et al., 2023). They had no history of neurological of psychiatric disorders, including auditory deficits, and were not taking any medication active in the central nervous system. All participants signed an informed consent form prior to participation. The study was approved by the regulatory and ethics administration in France (IDRCB nr. 2020-A01830-39).

### Task

Participants performed a passive auditory frequency oddball paradigm, which was a simplified version of the one used in (Lecaignard et al., 2015), while viewing a movie with audio muted unrelated to the stimulation, see Fig. 1A. One session of the auditory stimulation consisted of 2 blocks of 674 tones, totaling 1348 auditory tones, of which 1124 standards and 224 deviant tones (∼16.6%, i.e., 1/6). The tone onsets were spaced with a fixed interval of 610ms, tone duration was 70 ms including a 5 ms rise and fall (one session lasted approximately 14 minutes). Deviant tones differed in frequency from the standard tones (either 500Hz or 550Hz, which alternated across blocks, see Fig. 1A) and were intermixed randomly (uniform distribution) in the sequence, with at least 2 and at most 8 standard tones in between. This created an unpredictable sequency of tones. Stimuli were presented using Presentation software (Neurobehavioral Systems, Albany, CA, USA). Video was presented using a Propixx projector (VPIXX technologies Inc, Canada). Sounds were presented via a set of MEG-compatible earphones (EARTONE 3A, Etymotic Research Inc., 3m tube length), and adjusted to a comfortable level for the participant. Participants were instructed to ignore the sounds and focus on the (silent) movie.

**Figure 1.**
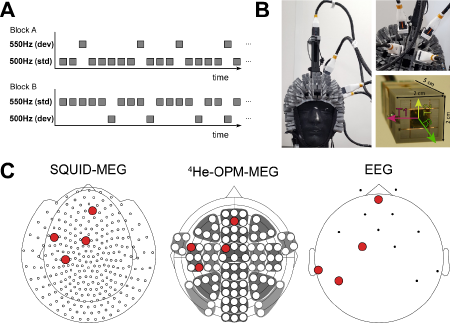
Experimental setup. A) Example timelines of the two blocks in a session. The frequency of the standard (std) and deviant (dev) stimuli was switched across blocks B) The OPM setup, with four sensors mounted in a semi-flexible helmet and one reference sensor mounted 10cm above the head. The scalp sensor locations in the photo are not the same as those used in the current study. A bottom view of the sensor is shown bottom right panel, with measurement axes (R: radial axis, T1, T2: tangential axes 1 & 2). The sensitive element (glass cell) can be seen in the center. C) Sensor layout for SQUID-MEG, OPM-MEG and EEG respectively, with sensors selected for comparison in red.

### Data acquisition

Participants completed one session with concurrent SQUID-MEG – EEG, followed by one session with concurrent ^4^He-OPM – EEG. We refer to the EEG session simultaneous with SQUID-MEG as ‘EEG_SQUID_’, and the EEG session simultaneous with the OPM-MEG acquisition as ‘EEG_OPM_’. The task was performed in a dimly lit magnetically shielded room (2 µ-metal, 1 copper layer, Vacuumschmelze, Hanau, Germany), while seated upright.

SQUID-MEG was recorded using a 275 axial-gradiometer MEG system (CTF MEG Neuro Innovations Inc., Port Coquitlam, BC, Canada). Concurrently, EEG (CTF EEG system, passive electrodes) was recorded from 14 scalp electrodes (see Fig. 1C), using the acquisition hardware integrated in the CTF-MEG system. A reference electrode was placed on the tip of the nose. Additionally, one bipolar, vertical EOG was recorded. Impedances were kept below 10 kOhm. Data were digitized at 1.2 kHz with a 300 Hz low pass-filter. In a separate session, concurrent OPM-EEG was recorded. The EEG setup was otherwise identical to the concurrent SQUID-MEG - EEG session. OPM-MEG recordings were performed using a set of five novel triaxial Helium-4 OPMs (MAG4HEALTH, Grenoble, France) (Beato et al., 2018; Labyt et al., 2019). The sensors were mounted in a semi-rigid helmet, carefully placed over the EEG. Four of the sensors were placed on the scalp (see Fig. 1B/C) and one sensor was mounted on a 10cm column on top of the head, serving as a reference sensor. Sensor positions were chosen to cover frontal and temporal areas, based on the known scalp topography of the MMN. Due to the low number of sensors, they were only placed on the left side. As the slots were open ended, the OPM sensors were placed as close to the scalp as possible. As these sensors operate at room temperature, there is no need for additional insulation between the scalp and the sensitive element of the OPM sensor. A wooden frame was used to support the cables from the OPM system and provide weight relief for the head. As these sensors have a large signal bandwidth (i.e. no saturation due to large fluctuations), no additional (active) shielding was used. The OPM-MEG data were digitized at 11 kHz.

### Data preprocessing

Data were analyzed using MNE-python (version 1.7.0., Gramfort et al., 2013) in a Linux environment. Data analyses of SQUID-MEG, OPM-MEG and EEG were aimed to be as similar as possible, while optimizing individual data quality. Despite EEG and MEG being recorded simultaneously, all modalities were processed separately, including artifact rejection, to be able to compare the end-results per modality. The EEG_SQUID_ and EEG_OPM_ sessions were also analyzed separately. The ^4^He-OPMs record from three axes simultaneously (one radial, two tangential axes), but the sensitivity on the second tangential axis is a factor 4 lower than the other axes (∼200 ft/√Hz vs. <45 ft/√Hz). Data from this axis was low pass filtered at 10Hz and used to identify artifactual components in independent component analysis (ICA), but not analyzed further and the analyses focused on the remaining radial and best tangential OPM axis.

First, for all modalities, segments in the data containing breaks (>9s), large amplitude fluctuations (>5 SDs from the median) and spikes (>5 SDs peak-to-peak) were marked to be omitted. Data were then bandpass filtered between 2 and 45 Hz.

Then, for the OPM data only, data from the reference sensor was used to de-noise the data. For this, the reference sensor data were low-pass filtered at 5Hz to avoid removing higher frequency content. These were then used to calculate a rolling regression over time (linear regression using a 2s sliding overlapping time window, 100ms steps) per channel, which was subsequently subtracted from the scalp sensor signal. The rolling regression was used to better adapt to head movements, which have a higher impact on the data than fluctuations in the background field.

Data from all modalities were then subjected to ICA (fastica, Hyvärinen and Oja, 2000) where components related to eye movement, muscle artefacts, blinks, heartbeat and gross non-neural artifacts (if present) were removed. The ability to isolate and remove these artefacts was more limited for OPM than the other modalities due to the low number of sensors and therefore ICA components (12 in total: four sensors with three axes). For SQUID-MEG 3.6 (±0.7) components were removed, for OPM-MEG this was also 3.6 (±0.8) components; for EEG_SQUID_ 3.2 (±1.3) components and EEG_OPM_ 4.4 (±1.1) components. The resulting data were subjected to a final 20 Hz low-pass filter and epoched relative to either the deviant tones, or the standard tone preceding the deviant [-.2 s to .41 s], which resulted in a maximum of 224 ‘standard’ and 224 ‘deviant epochs per subject. The epochs were then resampled to 1kHz and the EOG was used to regress out any residual eye movement artefacts. All EEG data were re-referenced to the average reference. Finally, epochs were baseline corrected using the 200ms up to the onset of the tone and any remaining artefactual trials were removed using the autoreject package (Jas et al., 2017). Final average reject rates were 0.5% (±1.1) / 0.8% (±1.3) for SQUID-MEG standard/deviant; 16.2% (±14.6) / 16.4% (±14.9) for OPM-MEG standard/deviant; 5.0% (±7.6) / 5.0 (±7.9) for EEG_OPM_ standard/deviant and 9.5% (± 9.8) / 9.1% (±9.6) for EEG_SQUID_ standard/deviant. Unaveraged individual rejection rates per modality can be found in Fig. 4C.

### Data analysis

For each modality, individual ERFs/ERPs for standard (preceding a deviant) and deviant tones were averaged over blocks to remove possible undesirable tonotopic effects. As the MMN is typically visible on the “deviant-minus-standard” response (hereafter called the difference response), we tested for significant differences between the standard and deviant ERFs/ERPs for SQUID-MEG, EEG_SQUID_ and EEG_OPM_ using spatiotemporal cluster permutation tests (Maris and Oostenveld, 2007). Due to the sparsity of EEG electrodes, the connectivity matrix for EEG had to be corrected by removing connections between distant electrodes, resulting in a decoupling of the left and right lateral electrodes (TP9/10 & P7/8) from the frontal and central electrodes. The cluster forming threshold was set at p = .01 and 10000 permutations were run. Due to the low number of sensors, OPM statistics were performed at the sensor level, per axis, on a time window selected based on the significant time interval for the SQUID-MEG data. Here, data from the standard and odd epochs were averaged in the selected time window [118 – 196 ms] per subject, sensor and axis and compared subject by subject using paired samples t-test. Results were Bonferroni corrected.

A second analysis was conducted to assess performance of the different modalities at the single subject level, given that clinical practice focuses on individual assessments. We used the same procedure as the group level statistical analysis, but at the single subject level across epochs. To directly compare SQUID-MEG, OPM-MEG, EEG_SQUID_ and EEG_OPM_, epochs were extracted and averaged for sensors with corresponding scalp location across modalities (see Fig. 1), based on Euclidean distance between sensor positions (SQUID: MLC25, MLC51, MLF46 & MRF41, EEG: AFz, C1, TP9 & P7). Sensor positions were based on the CTF sensor positions for SQUID-MEG, on the standard 1020 montage provided by MNE-python for EEG and on a digitized version of the helmet for OPM. Pearson product-moment correlations were calculated based on the averaged evoked time course [-200 – 410 ms] data at the group level. Furthermore, the signal-to-noise ratio (SNR) was calculated as the ratio of the maximum absolute MMN peak in a 100-250ms time window and the baseline standard deviation per sensor. To compensate any difference in the number of epochs, a sub selection of trials was used (equal to the least number of epochs) to equalize the number of trials. This was repeated 100 times to remove selection bias. As the MMN has its maximum at different sensors for different modalities, the sensor with the maximum SNR was chosen per modality.

## Results

We compared the performance of a novel ^4^He-OPM-MEG system with a ‘classic’ SQUID-MEG system and EEG in an auditory oddball paradigm. For every modality, typical auditory evoked responses could be measured by sensors/electrodes overlying the left temporal area (OPM sensor LT34 equivalent). ERF/ERPs from both standard and deviant stimuli show an early deflection between 61-74 ms on average (SQUID: 74ms, OPM_radial_: 71 ms, OPM_tangential_: 61 ms, EEG: 72 ms), followed by a deflection of opposite polarity between 111-166 ms (SQUID: 166 ms, OPM_radial_: 157 ms, OPM_tangential_: 111 ms, EEG: 139 ms).

### Significant MMN for SQUID-MEG, EEG and OPM-MEG

The grand average difference ERP/ERFs were highly similar across modalities, as can be seen in Figures 2 and 3 (see also supplementary figures 1-4). Permutation cluster test reveal significant mismatch negativities in all modalities, between approximately 100-200ms after tone onset. Specifically, SQUID-MEG shows both a left and right lateral cluster, with a maximum deflection (104 fT) between 118 and 196ms for the left cluster, and between 126ms and 202ms (82 fT) for the right lateral cluster. No other significant clusters were found. EEG results, recorded either simultaneously with SQUID-MEG or OPM-MEG, show an additional frontal cluster between 149ms and 202ms (.76 µV) for EEG_SQUID_ and similarly 150ms and 196ms (.70 µV) for EEG_OPM_. The left lateral cluster was found between 147 to 199 ms (1.57 µV) for EEG_SQUID_, but did not reach significance for the EEG_OPM_ session. Similarly, the right lateral cluster was significant for EEG_SQUID_ between 159 and 201 ms (1.19 µV), but failed to reach the significance level for EEG_OPM_ (p=0.061). OPM-MEG was analyzed per sensor and axis using the significant left cluster time window from the SQUID-MEG data (118-196ms). Here, both left lateral sensors’ radial axis shows a significant MMN (LT34 p<.001, 211 fT deflection amplitude; LT15 p= .015 corrected, -124 fT deflection amplitude). The tangential OPM axis also shows a weaker, but visible MMN (p< .05 uncorrected) in sensors LT34 and LC11, but these do not reach significance level after multiple comparisons correction (LT34 p= .40, -53 fT deflection amplitude, LC11 p= .14 corrected, -35 fT deflection amplitude). The frontal sensor did not show a significant deflection.

**Figure 2.**
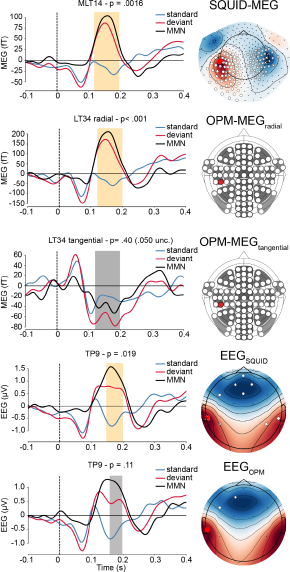
ERP/F statistics. Spatiotemporal clustering permutation test results for SQUID (top) and EEG (bottom). Significant clusters are indicated by enlarged symbols (circle, cross, diamond). For comparison, the time courses of the sensors with the strongest MMN (based on F score) of the left lateral cluster is plotted. For OPM, time courses are plotted for significant (based on time window analysis) sensor LT34. OPM p-values were Bonferroni corrected. For illustration purposes, selected non-significant results are included, indicated by grayed-out time windows. Full results can be found in supplementary figures 1-4.

**Figure 3.**
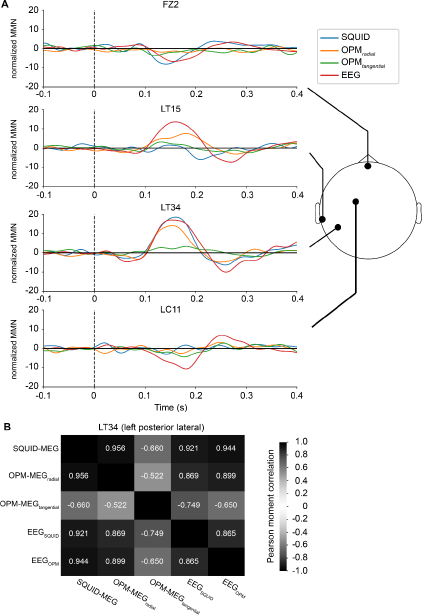
Time course comparison. A) Difference responses (deviant-standard ERF/ERP) for matching sensors. Colors indicate modality. The two EEG session are averaged for display purposes. B) Pearson moment correlations for the left posterior lateral sensor LT34 (or matching sensor/electrode for SQUID-MEG/EEG).

### High similarity for SQUID-MEG, OPM-MEG, EEG_SQUID_ and EEG_OPM_ difference ERF/ERP

A comparison of the difference ERF/ERPs from matching sensor locations can be seen in figure 3. As can be seen, the ERF/ERP time course is highly preserved across modalities, although the tangential OPM-MEG axis deviates most. This can also be seen in the time course correlations for the lateral sensors, where the radial axis shows a near perfect correspondence with the SQUID-MEG time course for posterior-lateral sensor LT34 (r=.96) and high correlation with the EEG data (EEG_SQUID_ r=.87, EEG_OPM_ r=.90). The tangential OPM-MEG axis also shows a good correspondence with SQUID-MEG (r=-.66) and EEG (EEG_SQUID_ r=-.75, EEG_OPM_ r=-.65), although the phase of the waveform is inverted. A similar pattern is observed for the fronto-lateral sensor LT15, with high correlations with EEG, phase inverted (EEG_SQUID_ r=-.91, EEG_OPM_ r=-.89), although SQUID-MEG deviates from the other modalities in this location. The tangential OPM-MEG axis mostly resembles EEG data (EEG_SQUID_ r=-.62, EEG_OPM_ r=-.70). Overall, this shows a high degree of reproducibility across the different modalities.

### Comparable performance for SQUID-MEG, OPM-MEG and EEG

Signal-to-noise ratios (SNR) were calculated based on the individual evoked MMN and results are displayed in figure 4. As can be seen, median SNR values are very similar across modalities, with SQUID-MEG having a larger number of high SNR individual values. Statistically, no difference was found between modalities in a repeated measure ANOVA (F_(4,64)_=2.12, p=.09). ERFs at the group level obtained using the OPM-MEG sensors thus seem to produce the same quality data as either SQUID-MEG or EEG.

**Figure 4.**
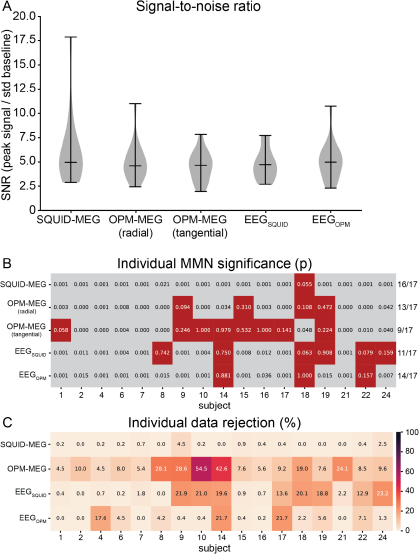
Performance comparison. A) Violin plots of signal-to-noise ratios for the modalities tested. The central horizontal black line denotes the median. B) Significance values per subject. P-values denote the minimal p-values obtained through either spatiotemporal clustering (EEG, SQUID-MEG) or time window analysis. Non-significant (p>.05) subjects are marked in red. C) Data percentage rejected per individual subject after preprocessing. Darker colors indicate more data rejected.

Looking at the within-subject statistics (Fig. 4B) it is clear than SQUID- MEG is more robust at the individual level, with 16 out of the 17 subjects showing a significant (p < .05) MMN in the expected time window. For the one subject where MMN was non-significant (S18, p=0.055), MMN was also non-significant using radial OPM-MEG, EEG_SQUID_ and EEG_OPM_. Using OPM-MEG, the radial axis data (OPM-MEG_radial_) yielded more individual significant MMN than the tangential axis (13 and 9, respectively) but less than SQUID-MEG. Data rejection rates (Fig. 4C) were generally higher for OPM. Regarding EEG, the two datasets (EEG_SQUID_, EEG_OPM_) differ slightly (11 and 14 individual MMN, resp; 14/17 similar conclusions). Interestingly, some participants that do not show a significant MMN with EEG do show one with OPM (even in a concurrent session; S08, S14, S18 and S22), and vice versa (S09, S15 and S19). While the tangential OPM axis generally performs poorer, it does detect significant modulations in a participant that does not show a significant modulation in other modalities (S18). Overall, the OPM-MEG, particularly the radial axis, performs on par with EEG (or better), but not SQUID-MEG. This is true despite higher rejection rates for OPM, resulting in a lower number of trials.

## Discussion

The aim of this study was to evaluate a new generation of MEG sensors, the ^4^He-OPMs, using human auditory evoked responses. Given the great potential of OPMs for routine clinical use, perhaps even at the patient’s bedside, which is not feasible using cryogenic MEG, we focused our analysis on the mismatch negativity (MMN): an auditory biomarker of perceptual and cognitive processes used in a wide range of neurological and psychiatric pathologies. Brain responses to tone sequences with oddball stimuli eliciting the MMN were recorded using magneto- encephalography (MEG) from SQUID and OPM sensors (in separate sessions) while participants watched a silent movie. At the time of data acquisition, we used a prototype OPM system consisting of only 5 sensors (including a reference sensor), that we positioned on the left frontotemporal part of the head surface. We also simultaneously recorded EEG during each session, as this is the standard clinical use of the MMN. We found typical MMN in all modalities and highly similar results across all modalities, showing promise for the novel ^4^He-OPMs to combine the advantage of MEG spatial precision with the flexibility of EEG.

The MMN is thought to originate from bilateral temporo-frontal generators (Fulham et al., 2014; Lecaignard et al., 2021; Rinne et al., 2000). Looking at the scalp topographic maps, a bilateral temporal dipolar pattern is usually dominant in the MEG signal (Hari et al., 1984; Rinne et al., 2000), while a central frontal deflection tends to be the strongest activity in EEG topographic maps, although this can depend on the reference (Giard et al., 1990; Kasai et al., 1999). Our current results are consistent with this. The results from SQUID-MEG indeed show strong bilateral temporal MMN clusters. A very similar result was obtained for the left hemisphere using the OPM sensors, despite there being only four scalp sensors. The radial axis posterior lateral OPM shows a remarkably similar time course to the ‘best’ left lateral SQUID-MEG sensor, but with about twice the amplitude (the MMN difference wave peaks at ∼211 fT for radial OPM, ∼104 fT for SQUID). Interestingly, the radial axis of the more frontal lateral OPM sensor shows a significant MMN with reversed polarity relative to the posterior lateral sensor, which did not reach significance for SQUID-MEG even though there is a clear lateral dipolar scalp pattern. This may be due to a placement of the OPM sensors closer to the scalp, resulting in better signal. With the rigid helmet employed in SQUID-MEG, the distance of the sensor to scalp can increase for more lateral sensors, depending on the participants head shape and size. Given that the dipole magnetic field follows an inverse square law with distance, this can be a significant change in signal power (Bonnet et al., 2025; Boto et al., 2016; Iivanainen et al., 2017). While the tangential axis of the posterior OPM sensors also shows the MMN, this did not survive statistical correction. Finally, the EEG results, from both sessions (concurrent with SQUID-MEG and OPM-MEG) show a significant frontal MMN.

EEG can be seen as complementary to MEG, as MEG is insensitive to (perfectly) radially oriented sources, while EEG is relatively invariant to source orientation (Ahlfors et al., 2010). Unfortunately, this is not ameliorated by the sensitivity profile of the OPM axes (radial, tangential), as this ‘radial blindness’ is intrinsic to the magnetic signal. Where radial MEG sensors, such as most SQUIDs and radial axis OPM are sensitive mostly to primary currents, tangential sensors are more sensitive to volume currents, as is EEG (Iivanainen et al., 2017). In a recent study using the same OPM system measuring epileptic activity, the tangential OPM axis provided a delayed component that may be informative on the propagation of interictal activity (Badier et al., 2023).

While the tangential axes may record additional brain activity, it is currently difficult to interpret. It is clear that there is real advantage to the additional axis in noise reduction. The different axes of the OPM sensor add additional ‘viewpoints’, allowing for a better characterization of the magnetic field in the presence of external noise (which is usually the case). This enables a better separation of noise and brain signal (Brookes et al., 2021; Tierney et al., 2022). This advantage is not apparent in the current study due to the low number of sensors, prohibiting the use of advanced noise suppression techniques (HFC, (Tierney et al., 2021), AMM (Tierney et al., 2024) or source localization (i.e. beamformers Gross et al., 2001).

Although recorded from the same participants in separate sessions, the EEG results showed some differences between the sessions concurrent with SQUID-MEG and OPM-MEG. The EEG session concurrent with the OPM recording showed only a single (frontal) cluster for the MMN, while the session concurrent with SQUID-MEG showed three significant clusters. In contrast, the OPM-EEG session shows more individually significant results. However, we do not believe this points to a relevant difference. The results are more similar than they are different when comparing the group time courses. It is likely that there is a slightly higher variability in the OPM-EEG session, which is also reflected in the wider spread of SNR values. This may have to do with the OPM helmet fitting over the EEG sensor, potentially causing a variability in recording quality. Furthermore, due to practical restrictions the EEG session concurrent with SQUID-MEG was performed first, which may have led to a difference in fatigue levels.

When directly comparing the difference response obtained from SQUID-MEG, OPM-MEG and EEG, the group average time courses are strikingly similar. The radial axis OPM ERF has high time course correlations (r >.87) with all other modalities, indicating that the OPM system was able to capture the same neural dynamics with high precision equivalent to SQUID-MEG and EEG. The tangential OPM axis shows a clearly deviating pattern of activity from all other time courses, which is not unexpected given its different sensitivity profile. The sensor placement was optimized in line with the known scalp patterns from EEG and SQUID-MEG (Lecaignard et al., 2021). The tangential axis therefore, being sensitive to a different aspect of the magnetic field than the radially sensitive SQUID sensors, may not have been in the optimal location. Furthermore, the amplitude of the tangential component has an inherently lower amplitude than the radial axis (Iivanainen et al., 2017). Despite this, the signal-to-noise ratio is comparable to the other sources. Importantly, in addition to the shape of the time course, the SNR is near equivalent across the board, with only a slight advantage for SQUID-MEG.

At the individual level, the OPM system also shows good performance in detecting MMN on par with EEG. This is especially relevant for clinical applications, where the MMN is used as a biomarker. While the fully developed SQUID-MEG system has the superior performance here, it must be noted that this analysis was performed using all available sensors, also for EEG, which is a clear benefit in both preprocessing and sensor-level analysis. Thus, despite this disadvantage, the good performance of the novel OPM-MEG system is promising for individual diagnose.

While the OPM-MEG system has shown near equivalent performance, the system used is comprised of only four scalp sensors and a single reference sensor. This study is part of the first steps of a real-world evaluation of the Helium OPM sensors, and suffers from limitations of a prototype system, most prominently the low number of sensors. The experiment took place in a magnetically shielded room, but without any active shielding. These active compensation coils, aimed at further nulling the background field, are becoming increasingly common in use with lower dynamic range alkali OPM-MEG (Hill et al., 2022). Due to the large dynamic range of the ^4^He-OPMs the recording session was not affected by large magnetic field fluctuations, but the recorded data do contain large artefacts due to subject movement and changing external magnetic fields. Unlike SQUID-MEG, which uses gradiometers that natively attenuate distant noise sources, the (OP) magnetometers record all of the magnetic field, near and far. We used a relatively simple preprocessing pipeline based on reference regression, ICA and filtering. As mentioned above, the low number of sensors puts severe limitations on the ability to use more effective noise reduction techniques. This is reflected in higher data rejection rates compared to SQUID-MEG and EEG.

However, at time of writing, a new iteration of the ^4^He-OPM has been developed with a lower noise floor better than 30 fT/√Hz (as compared to the ∼45 fT/√Hz used in the current study). Future steps involve testing this system with improved noise floor and increased number of sensors, providing whole head coverage. This will enable more effective post-processing, as well as source modeling. For now, given all the mentioned limitations, the on-par performance of the system holds promise for high-quality, wearable MEG that combines the practical advantages of EEG and signal quality of MEG, creating potential for a flexible, high-quality setup for routine clinical use.

## Acknowledgements

This project was funded by Région Auvergne-Rhône-Alpes (Pack Ambition project: NEW_MEG). This project was partly funded by France Life Imaging (WP2 & 4) (grant “Infras- tructure d’avenir en Biologie Santé” ANR-11-INBS- 0006). TG is supported by the Labex Cortex (ANR-11-LABX-0042).

## Data availability statement

Anonymized data supporting the results of this study are available from the corresponding author upon reasonable request and validation by regulatory and ethical bodies and subject consent. Scripts used for analysis can be found at https://github.com/tgutteling/OPM_MMN

## Ethics statement

The study was approved by the regulatory and ethics administration in France (IDRCB nr. 2020-A01830-39).

