## supplementary figures for "The Mismatch Negativity compared: EEG, SQUID-MEG and novel ^4^Helium-OPMs"

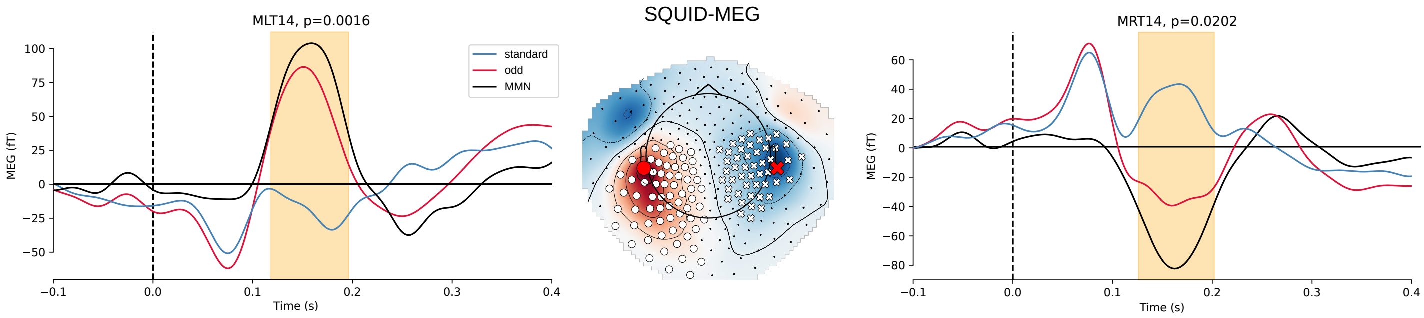


*Supplementary figure 1: Full spatiotemporal clustering permutation tests results for SQUID-MEG. Significant clusters are indicated by enlarged symbol (circle, cross). For each cluster, the timecourse of the sensor with the strongest MMN (based on F score) is plotted. Significant time window is indicated in yellow.*

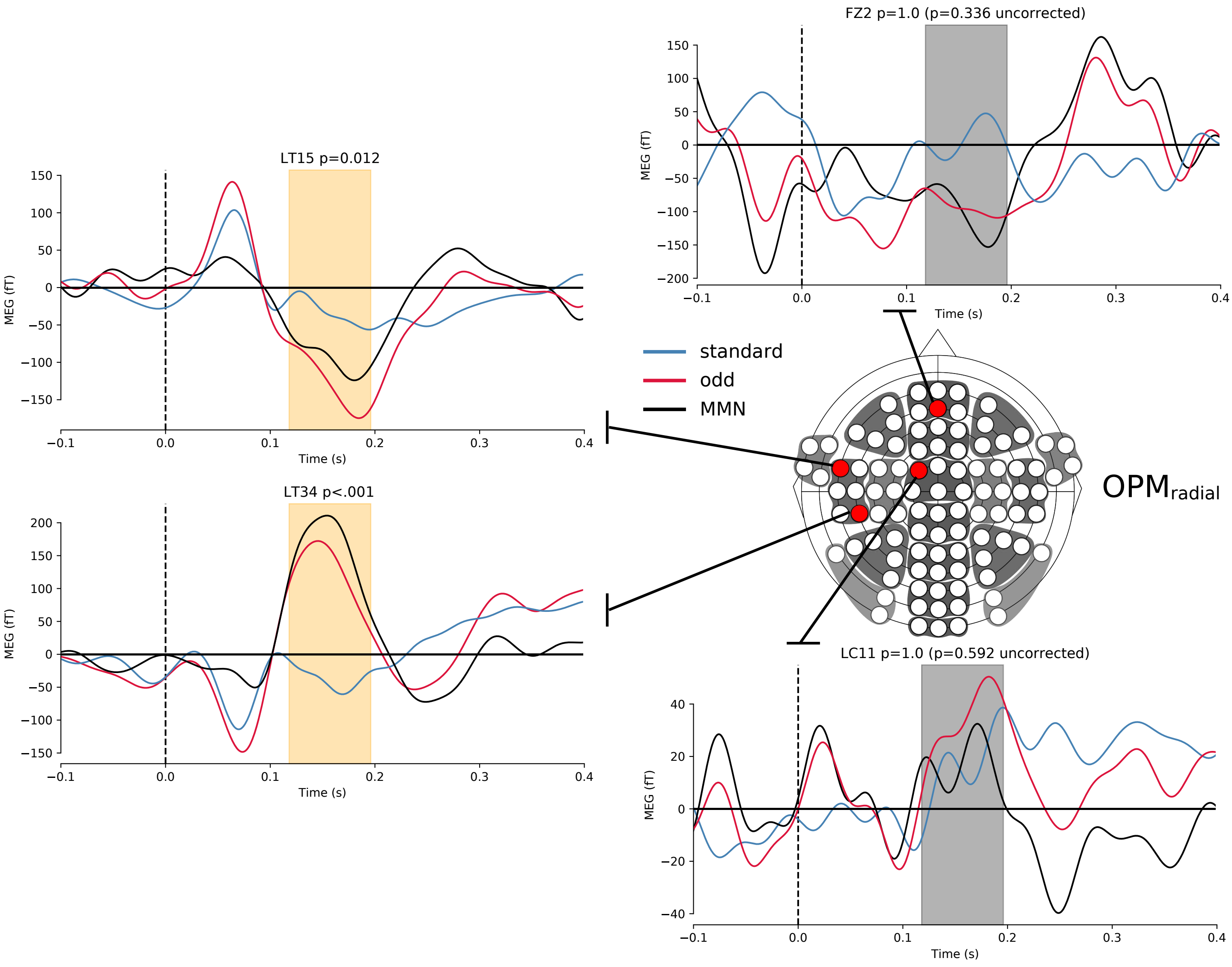


*Supplementary figure 2: Full time window statistics for the OPM-MEG$_{radial}$ data. Time courses are plotted for all sensors (radial axis only). OPM p-values were Bonferroni corrected. Significant results are marked in yellow, non-significant results are indicated by grayed-out time windows.*

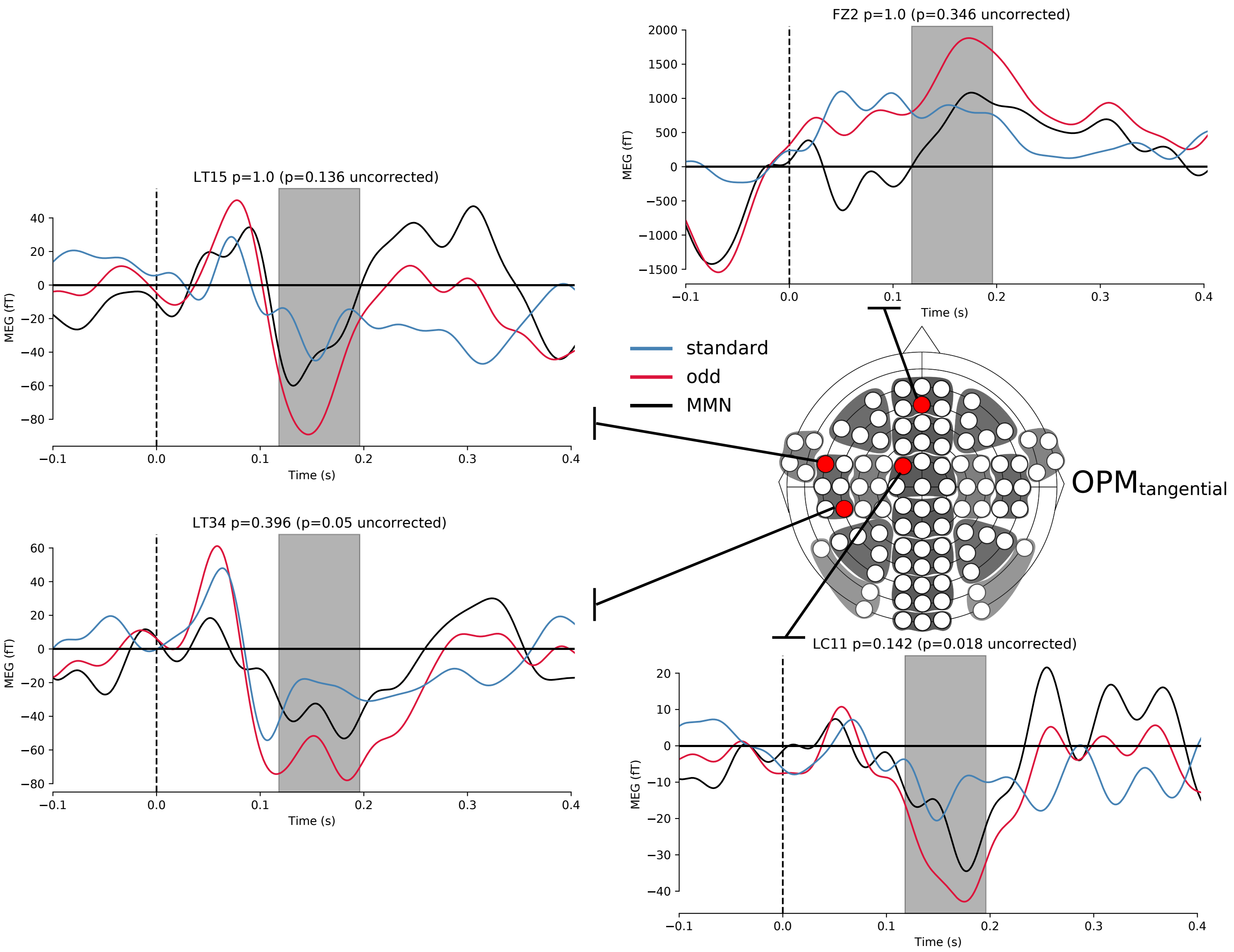


*Supplementary figure 3: Full time window statistics for the OPM-MEG$_{tangential}$ data. Time courses are plotted for all sensors (tangential axis only). OPM p-values were Bonferroni corrected. Nonsignificant results are indicated by grayed-out time windows.*

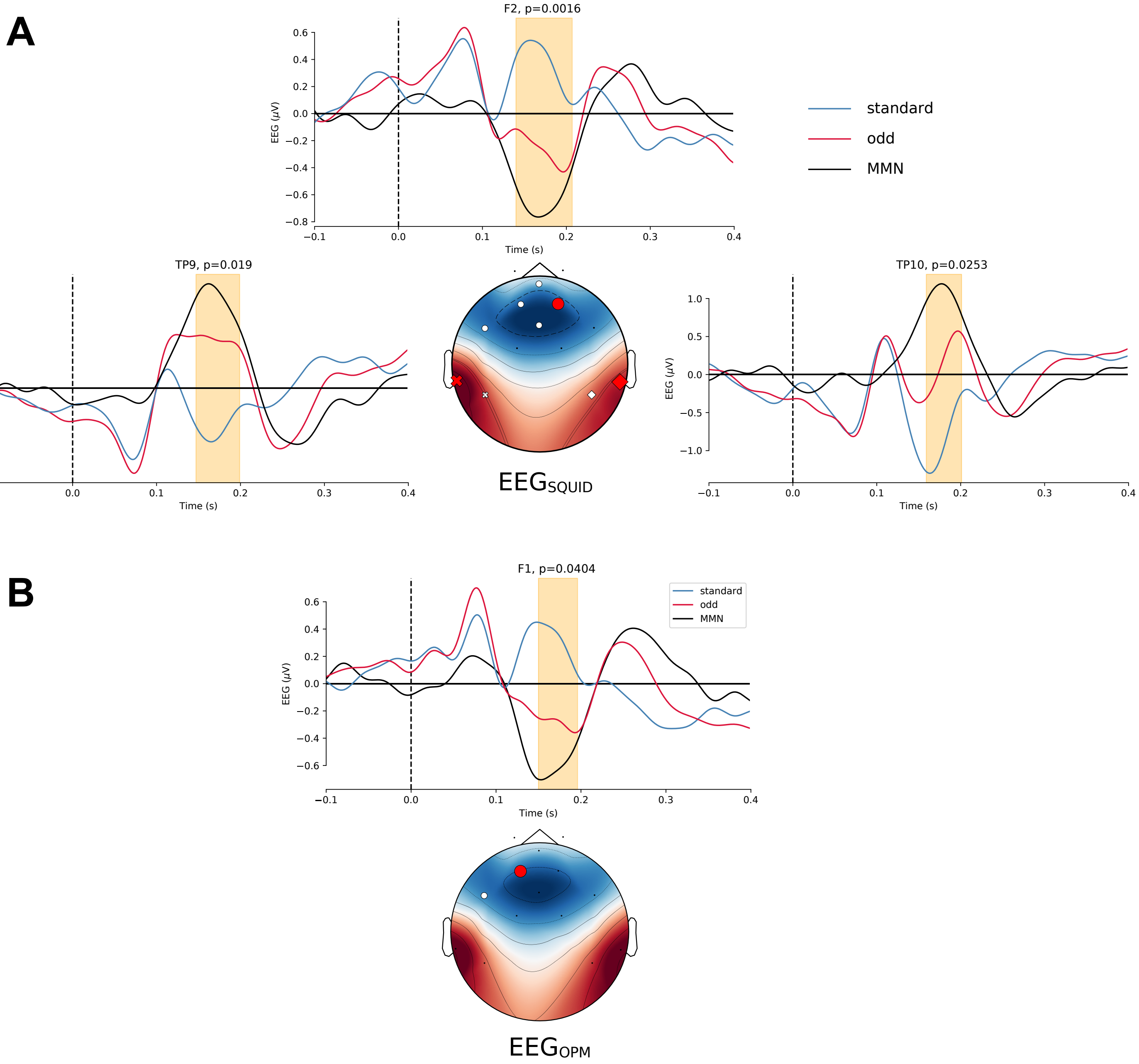


*Supplementary figure 4: Full spatiotemporal clustering permutation tests results for the EEG data, for the EEG session concurrent with SQUID-MEG (A) and concurrent with OPM (B). Significant clusters are indicated by enlarged symbol (circle, cross, diamond). For each cluster, the timecourse of the sensor with the strongest MMN (based on F score) is plotted. Significant time window is indicated in yellow.*
